# Single Pulse Heating of Nanoparticle Array for Biological Applications

**DOI:** 10.1101/2021.10.21.465356

**Authors:** Chen Xie, Peiyuan Kang, Johan Cazals, Omar Morales Castelán, Jaona Randrianalisoa, Zhenpeng Qin

## Abstract

With the ability to convert external excitation into heat, nanomaterials play an essential role in many biomedical applications. Two modes of nanoparticle (NP) array heating, nanoscale-confined heating (NCH) and macroscale-collective heating (MCH), have been found and extensively studied. Despite this, the resulting biological response at protein level remains elusive. In this study, we developed a computational model to systematically investigate the single-pulsed heating of NP array and corresponding protein denaturation/activation. We found that NCH may lead to targeted protein denaturation, however, nanoparticle heating does not lead to nanoscale selective TRPV1 channel activation. The excitation duration and NP concentration are primary factors that determine a window for targeted protein denaturation, and together with heating power, we defined quantified boundaries for targeted protein denaturation. Our results boost our understandings in the NCH and MCH under realistic physical constraints and provide a robust guidance to customize biomedical platforms with desired NP heating.

## 1. Introduction

Nanomaterials can be designed to efficiently absorb optical or magnetic energy and convert that energy to heat.^[1–3]^ This heat-generating property has found applications in cancer treatment,^[4–8]^ drug delivery,^[9, 10]^ contrast imaging,^[11–14]^ vision restoration,^[15]^, pain^[16]^ and neuromodulation (Figure S1).^[17, 18]^ While these advances have pushed the boundaries of biomedicine, there has been a fair amount of debate over the past few years in understanding the effect of heating in biological systems, such as in the area of magnetogenetics on whether magnetothermal heating of ferritin nanoparticles can actually generate enough heat to actuate thermally-sensitive ion channels.^[19–23]^ Scaling analysis suggests that the heat generated by ferritin is 9-10 order of magnitude less than that threshold to activate TRPV1 ion channels^[24]^ and recent reports suggest a biochemical pathway involving reactive oxygen species (ROS) that activates both TRPV1 and TRPV4 ion channels.^[23, 25]^ This highlights the importance of understanding nanoscale heating and its impact on biological systems, especially proteins.

Heating generated by nanomaterials broadly falls into one of two regimes: nanoscale-confined heating (NCH) and macroscale-collective heating (MCH). NCH is characterized by local temperature increases at the nanoscale which remain confined near the surface of the nanoparticle, as in the case of controlling protein activity by molecular hyperthermia,^[15, 16, 26]^ or membrane permeability by opto-poration.^[16, 26–28]^ In contrast, MCH is characterized by a widespread temperature increase throughout the medium, such as whole cells or tissues. MCH can, for example, be used to injure cancer cells,^[5]^ excite thermally-sensitive neurons expressing transient receptor potential (TRP) channels,^[15]^ and regulate cellular function.^[29]^ Keblinski *et al.* highlighted the role of the duration of energy excitation in controlling the transition between NCH and MCH,^[30]^ demonstrating that short energy pulses could lead to NCH while continuous excitation could cause MCH. Similarly, Baffou *et al.* systemically investigated the boundary and transition between temperature confinement (or NCH) and delocalization (or MCH) under continuous excitation and repeated femtosecond pulsed excitations, respectively.^[31, 32]^ Furthermore, methods for generating arbitrary microscale temperature profiles have been explored using NPs and spatial light modulation.^[33]^ While these efforts advanced our understanding of the spatiotemporal evolution of nanoparticle heating, there is an important gap in understanding how single pulsed NP heating affects biological systems through protein denaturation (by hyperthermia^[5]^ or molecular hyperthermia^[16]^) or activation (for example thermally sensitive ion channels TRPV1).^[15]^

In this report, we investigated the heating of NP arrays under single pulse excitation and the resulting protein responses, including protein thermal denaturation and activation of thermally-sensitive ion channels. Our analysis suggests that the excitation duration and NP area density determine a window for heterogeneous protein denaturation, where targeted denaturation is possible. Together with heating power, we defined boundaries for targeted protein denaturation inside this heterogeneous denaturation window. For activation of thermally sensitive ion channel TRPV1, we determined that the nanoparticle array heating leads to widespread of channel activation instead of localized activation, because heat dissipates and leads to MCH within milliseconds, a timescale required to activate the ion channel based on our current understanding.^[34]^ This work provides guidance for designing innovative approaches that utilize NCH and MCH for modulating molecular or tissue-specific activities under realistic physical constraints.

## 2. Results and Discussions

### 2.1 NP array heating characteristics and corresponding protein denaturation

We first constructed and validated the modeling framework for NP array heating and protein thermal denaturation (Figure 1 A). For our model systems, we used 10 μm ×10 μm 2D square arrays composed of 30 nm gold nanoparticles (AuNP) immersed in water, which were designed to mimic NPs that are targeted to a cell membrane. The NP array heating was acquired by a superposition method (Equation S1). Our results suggest that different NP materials with the same heat generation will have a similar heating profile in water (Figure S2 A), which is the main target for analyzing heating in biological tissues. Therefore, we used heating power per NP (*g*) that is independent of NP absorption properties to represent the heat generation of NPs by different methods, such as laser or magnetic field (Figure S1). This heating model was used throughout the remaining of the paper.

**Figure 1.**
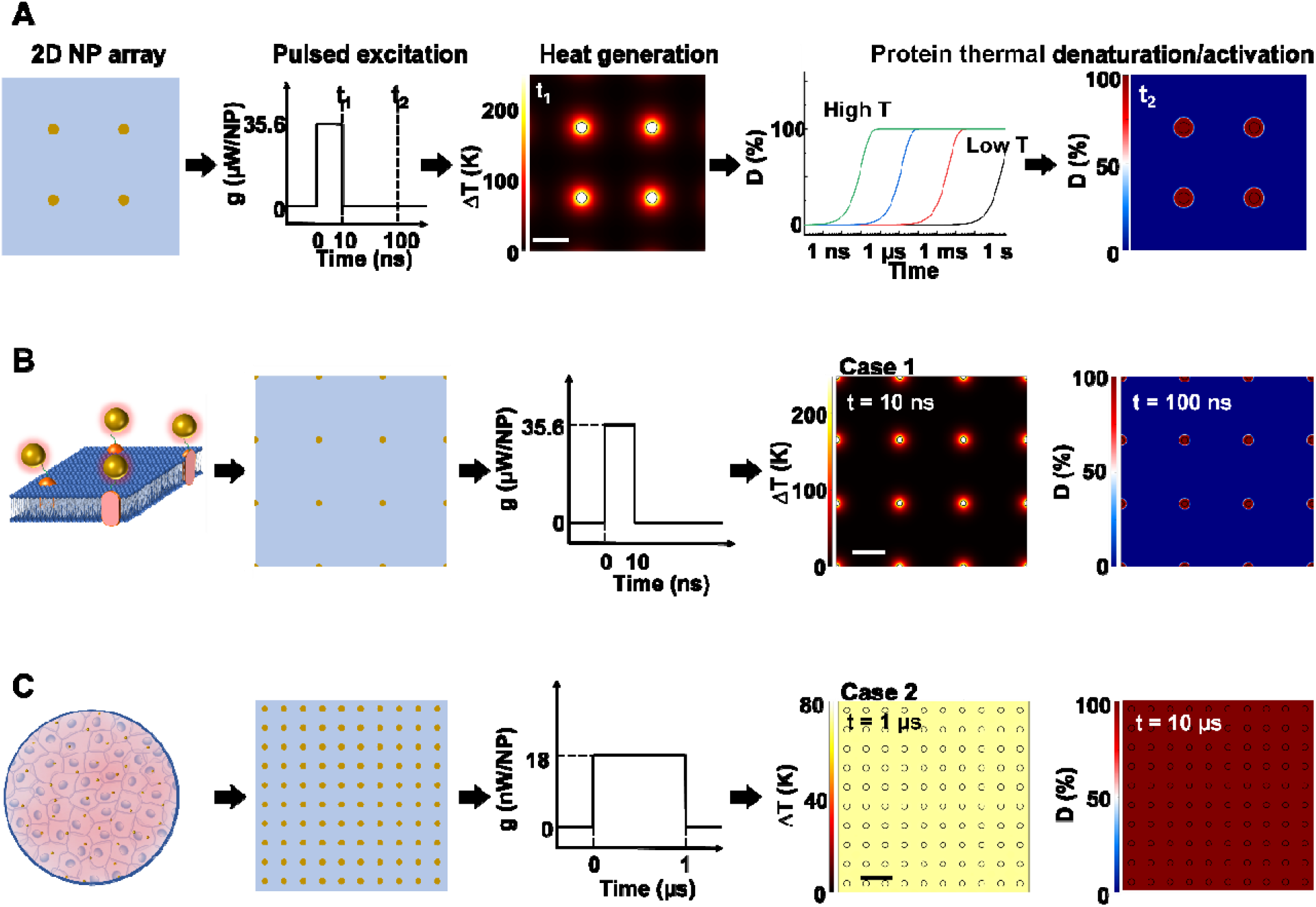
Analysis framework for nanoparticle heating-induced protein denaturation and characteristics of protein denaturation under nanoscale-confined heating (NCH) and macroscale-collective heating (MCH). (A) Schematic of the analysis framework for nanoparticle (NP) array heating and the resulting protein denaturation (*D*) or activation. External excitation of the NP array (each NP as a nano heater) heats up the NP and the surrounding media. The temperature increase drives a chemical reaction (protein denaturation or activation), which can be described by first-order kinetic model (the Arrhenius model). Integration of the reaction rate over time leads to an overall protein denaturation or activation. (B-C) Nanoscale confined heating (NCH) and macroscale collective heating (MCH), and the corresponding temperatur increase and protein denaturation. Conditions: domain size 10 μm ×10 μm, NP diameter (*d_NP_*) = 30 nm, heating power per NP (*g*) is (B) 35.6 μW and (C) 18 nW, NP area density is (B) 9 μm^−2^ and (C) 100 μm^−2^, and excitation duration is (B) 10 ns and (C) 1 μs.

Heating can trigger a series of biological responses, including protein denaturation in the case of cancer thermal therapy,^[6, 35]^ and activation of temperature-sensitive TRPV1 channels that modulate neuron activity.^[15, 36]^ Recently, Kang *et al.* showed the possibility of nanoscale selective protein denaturation with AuNP targeting and nanosecond pulsed laser excitation.^[16]^ To connect NP heating to the protein denaturation, we first analyzed protein denaturation using α-chymotrypsin since its denaturation properties have been measured in both NCH and MCH.^[28,37]^ Protein thermal denaturation can be modeled as a temperature-sensitive chemical reaction (Equation S3, Figure 1 A), and the normalized concentration of the denatured protein (*D*) is determined by the reaction rate and reaction duration (Equation S4). Since temperature profile and protein denaturation are all time dependent, we consider the end of excitation duration as the representative time point (*t*) for temperature change (Δ*T*) analysis, and the time after the laser pulse (10 × excitation duration) as the representative time point for protein denaturation analysis.

Similarly, we further consider the NP-water interface (P_1_) and the mid-point between NPs (P_2_) as two representative locations for Δ*T* and denaturation analysis (Figure S2 B). We will use P_1_ and P_2_ to indicate these locations, along with Δ*T_1_*, Δ*T_2_* and *D_1_, D_2_* to indicate temperature increase (Δ*T*) and normalized protein denaturation (*D*) at P_1_ and P_2_ respectively. These representative time points and locations are used throughout the paper. With this framework, we examined the characteristics of nanoscale-confined heating (NCH) and macroscale-collective heating (MCH) and connected them to corresponding protein denaturation. The thermal activation of TRPV1 was analyzed using a similar framework and presented in later sections. Figure 1 B shows that NCH was observed under short excitation duration (10 ns) and low NP area density (9 μm^−2^), where heating is confined to the nanoparticle surface (Figure S3 A-B). As a result, targeted protein denaturation is established where the protein denaturation was confined to the vicinity of the NPs (Figure1 B, Figure S3 A-B). By increasing the heating duration to 1 μs and the NP area density to 100 μm^−2^ (Figure 1 C), the heating fully dissipates and leads to MCH where the temperature increased uniformly. In this case, widespread protein denaturation was observed. Together, these two cases demonstrate the major hallmarks of NCH and MCH and the resulting targeted *versus* widespread protein denaturation.

### 2.2. Key factors that affect the NP array heating modes and protein denaturation

Next, we identified the key factors that impact the transition from NCH to MCH and from targeted to widespread protein denaturation. We first investigated the role of excitation duration by comparing the thermal responses and corresponding protein denaturation of an NP array (area density of 9 μm^−2^) at three distinctive excitation durations (Figure 2 A; case 3: 10 ns, case 4: 100 ns, and case 5: 1 μs; all under same heating power (*g* = 35.6 μW); summary of all cases in Table S3). Here we adopt Δ*T_2_*/Δ*T_1_* and *D_2_/D_1_* to characterize the temperature change and denaturation profile. High Δ*T_2_/*Δ*T_1_* and *D_2_/D_1_* indicate uniform Δ*T* and denaturation profiles, while low Δ*T_2_/*Δ*T_1_* and *D_2_/D_1_* indicate heterogeneous profiles. Figure 2 B-D illustrate the Δ*T* profiles and protein denaturation for these cases. For case 3 (10 ns), heating was confined and Δ*T_2_* remained constant (Figure 2 B&D), which resulted in low Δ*T_2_/*Δ*T_1_* (Figure 2 E); while for cases 4 and 5 (100 ns and 1 μs), Δ*T_2_* increases significantly, leading to an increase of Δ*T_2_/*Δ*T_1_*. Cases 3 & 4 (10 and 100 ns excitation) resulted in targeted denaturation with low *D_total_* (1.6% and 4.8% respectively, Equation S5) and *D_2_/D_1_* (~ 0 %). In contrast, Case 5 (1 μs) exhibited widespread protein denaturation (*D_total_* = 99%, *D_2_/D_1_* = 100 %). In this scenario, we observed a sharp increase in Δ*T_2_/*Δ*T_1_*, *D_2_/D_1_* and *D_total_* when the excitation duration increases (Case 5, Figure 2 E). Therefore, the excitation duration is considered as a key factor in both transitions (*i.e.,* from NCH to MCH and from targeted to widespread denaturation).

**Figure 2.**
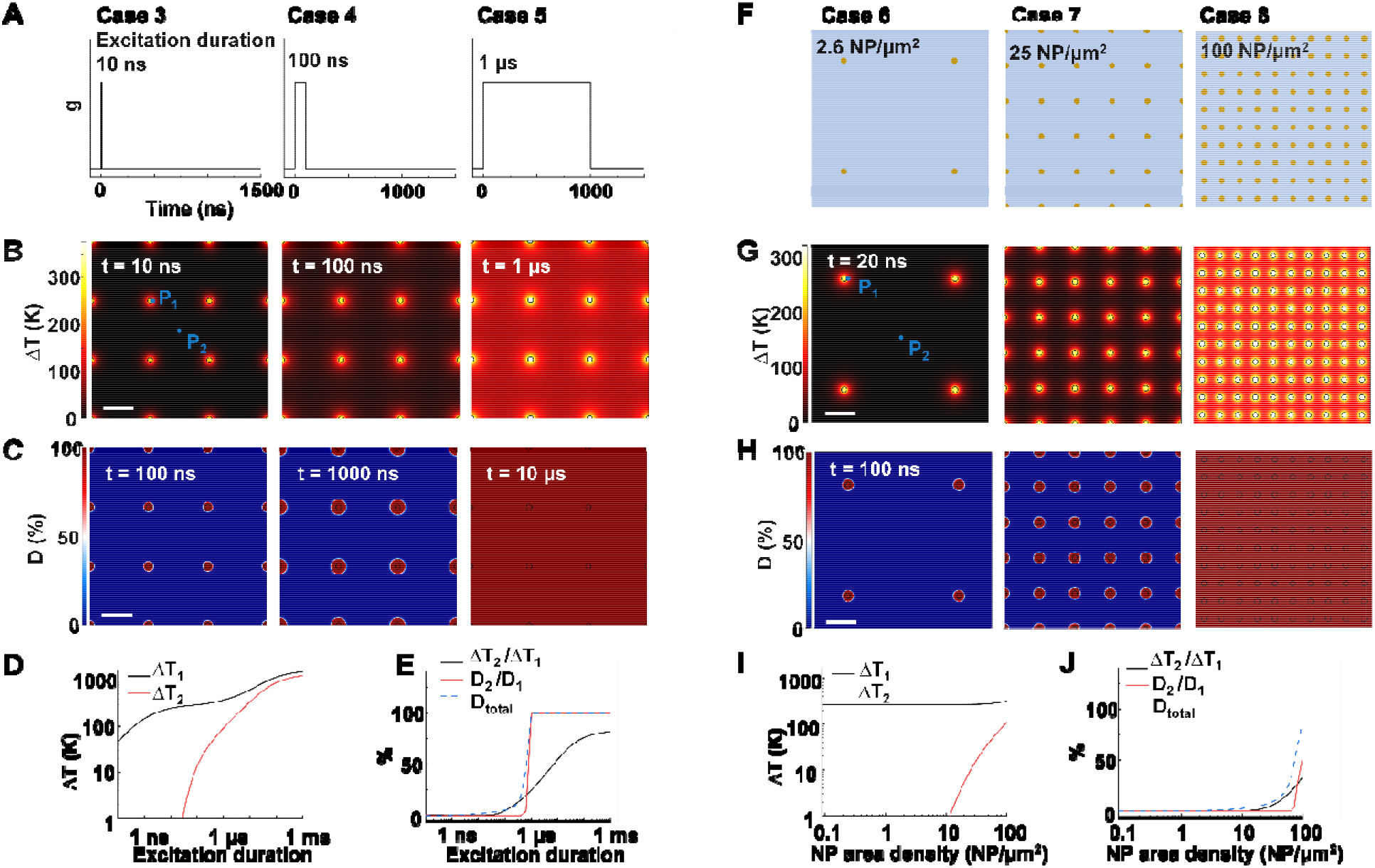
Effects of excitation duration and NP area density on the heating and protein denaturation. (A) Schematic of excitation durations for case 3-5. (B) Δ*T* profile and (C) protein denaturation (*D*) profile for cases 3-5. For case 3-5, NP area density is at 9 μm^−2^. P_1_ represents the NP-water interface, P_2_ represents mid-point between NPs. (D) Δ*T* at P_1_ (Δ*T_1_*) and Δ*T* at P_2_ (Δ*T_2_*) as a function of excitation duration. (E) Ratio between Δ*T_2_* and Δ*T_1_* (Δ*T_2_*/Δ*T_1_*), ratio between protein denaturation at P_2_ and P_1_ (*D_2_/D_1_*), *D_total_* in terms of excitation duration. (F) Schematic of NP area density for cases 6-8 at 2.6 μm^−2^, 25 μm^−2^, and 100 μm^−2^, respectively. (G) Δ*T* profile and (H) *D* profile for cases 6-8. For cases 6-8, excitation duration 20 ns. (I) Δ*T_1_* and Δ*T_2_* in terms of NP area density. (J) Δ*T_2_/*Δ*T_1_*, *D_2_/D_1_* and *D_total_* in terms of NP area density. For all cases, *g* = 35.6 μW and scalebar represents 200 nm.

When targeting and manipulating protein receptors on the cell surface, the NP area density depends on the receptor density. For example, metabotropic glutamate receptors (mGluRs) have an area density of 2 ~ 51 μm^−2^,^[2,13]^ requiring a similar NP area density on the cell surface for effective targeting. To evaluate the role of NP area density, we compared thermal responses and protein denaturation under three different NP arrays with distinct NP area densities (Figure 2 F; case 6, 2.6 μm^−2^; case 7, 25 μm^−2^; case 8, 100 μm^−2^; all under identical excitation condition: excitation duration 20 ns, *g* = 35.6 μW). Figure 2 G-H shows that, at low and medium area densities (2.6 μm^−2^ and 25 μm^−2^), NCH and targeted protein denaturation is observed. In contrast, at high area density (100 μm^−2^), heating generated by adjacent NPs are overlapped, resulting in MCH (Δ*T_2_/*Δ*T_1_* = 34.2 %) and widespread denaturation (*D_total_* = 100%, *D_2_/D_1_* = 100 %). Based on this analysis, we consider the NP area density as another key factor in both transitions.

The spatial distribution of NPs on the cell membrane is dependent on the distribution of target receptors, which can be randomly distributed.^[13, 38]^ To evaluate the role of NP distribution, we examined thermal responses and protein denaturation for three different NP arrays with identical area density (9 μm^−2^) distributed in square (case 9), hexagonal (case 10) and random (case 11) patterns (Figure S4 A). When subjected to a 20 ns excitation at 35.6 μW/NP, all of the NP arrays exhibited NCH and targeted protein denaturation (Figure S4 B&C). However, a close examination revealed that random distribution causes local temperature hot spots at the regions with NP aggregations (Figure S4 and Figure S5). This agrees with previous reports which showed that local temperature can be much higher around NP clusters.^[39, 40]^ It worth pointing out that *D_total_* of all three cases are similar due to the few local hot spots, making the NP distribution less significant for protein denaturation.

Next, we analyzed the effect of heating power (*g*) on the protein denaturation. Δ*T* increased linearly with *g* (Table S 1-2), resulting in a similar Δ*T* profile with different magnitude. On the other hand, the corresponding denaturation increased nonlinearly as per the Arrhenius equation (Equation S3). Here we compared cases under different heating powers with identical excitation duration (1 μs) and NP area density (9 μm^−2^). We identified three stages (I-III) of protein denaturation based on how *D_2_/D_1_* changes with the *g*. Figure 3 A and Figure S6 show that, in Stage I (homogeneous intact stage, case 12 as an example), there is no significant protein denaturation throughout the system due to insufficient *g* (both *D_2_* and *D_1_* are small, leading to high *D_2_/D_1_* ratio). In the Stage II, *g* is sufficient to generate a localized hot area and significant denaturation around NPs (case 13 as an example), with no significant heating or protein denaturation at P_2_ (*D_2_* ~ 0%). Therefore, a heterogeneous denaturation profile was observed with a small *D_2_/D_1_* ratio (< 1%). In the Stage III (case 14 as an example), *g* is sufficient to heat up the whole system and denature all proteins regardless of position, leading to homogeneous denaturation. We thus define two critical heating powers, *g_1_* and *g_2_* at *D_2_/D_1_* = 1% as boundaries for heterogeneous denaturation (Figure 3 A). It is worth noting that *g_1_* and *g_2_* are dependent on excitation duration and NP area density (Figure 3 B). Stage II vanishes at high NP area density and excitation duration (Figure 3 B, 100 μm^−2^, 1 ms), with a direct transition from no denaturation to widespread denaturation. This agrees with previous sections, where long excitation duration and high NP area density lead to MCH (high Δ*T_2_/*Δ*T_1_*). As such, there is no window for heterogeneous or targeted denaturation. To quantify the threshold for sufficient denaturation at P_1_, we further define a heating power *g_3_* corresponding to *D_1_* = 50% (Figure 3 C).

**Figure 3.**
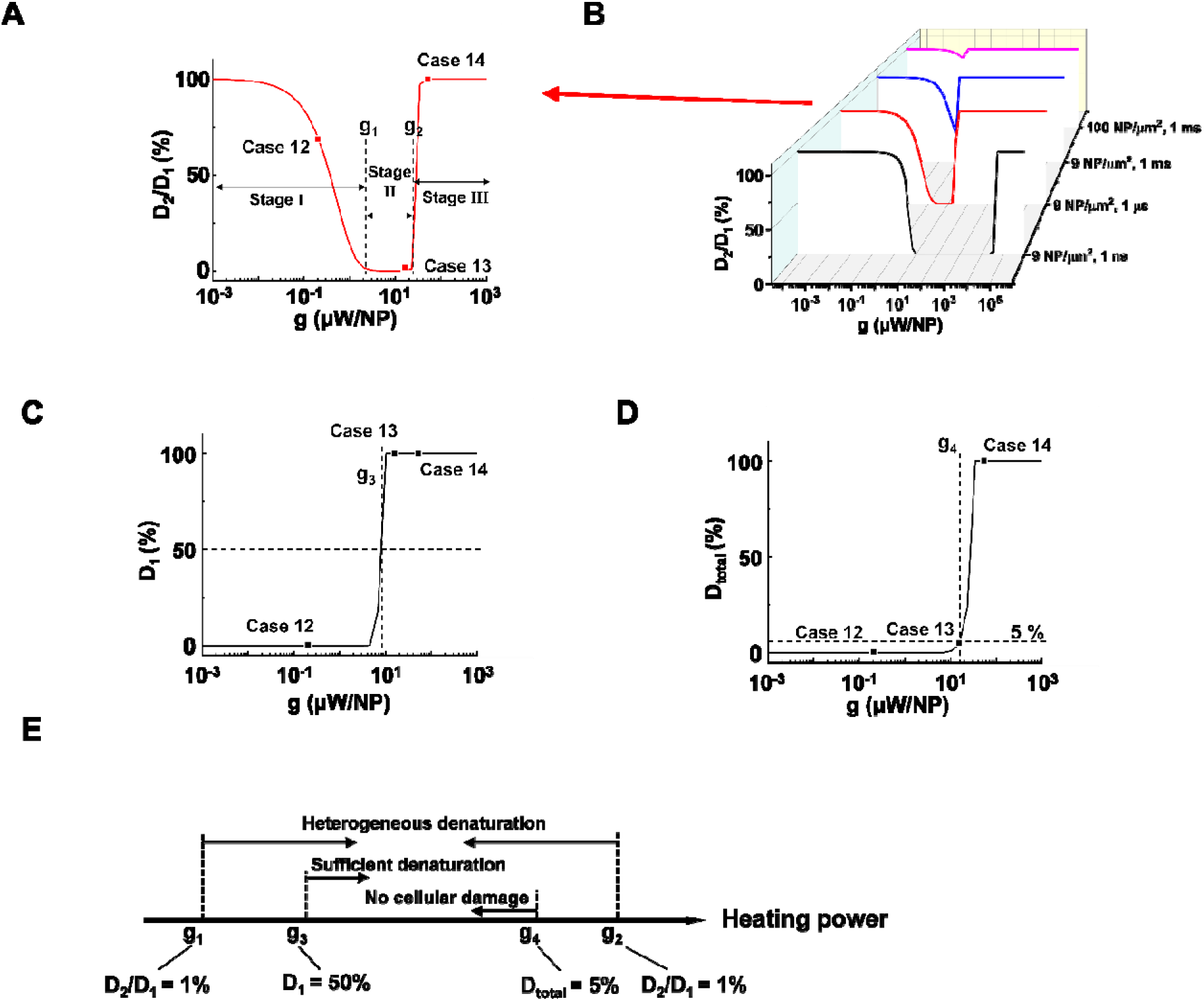
NP heating power determines a window for targeted denaturation. (A) *D_2_/D_1_* in terms heating power (*g*) with a 1 μs excitation duration and a 9 NP/μm^2^ area density. Three major stages for protein denaturation as *g* increases: Stage ◻, uniformly no denaturation zone with high *D_2_/D_1_* (1% – 100%); Stage ◻, targeted denaturation window with low *D_2_/D_1_* (< 1%); Stage ◻, widespread denaturation with high *D_2_/D_1_* (1% – 100%). *g_1_* and *g_2_* are critical heating powers that separate the three stages. (B) *D_2_/D_1_* in terms of *g* with different combinations of excitation durations and NP area densities. (C) *D_1_* in terms *g* with a 1 μs excitation duration and a 9 μm^−2^ area density. Here we define a critical heating power *g_3_* that corresponds *D_1_* = 50 %. (D) *D_total_* in terms *g* with a 1 μs excitation duration and a 9 μm^−2^ area density. We define a critical heating power *g_4_* that corresponds *D_total_* = 5 %. (E) Schematic for *g_1_*, *g_2_*, *g_3_*, and *g_4_*.

Heating power that is much lower than *g_3_* does not produce sufficient denaturation around NPs. Furthermore, studies have suggested a large total protein denaturation can lead to cellular damage (e.g. *D_total_* = 5 %) ^41^. Therefore, we define critical heating power *g_4_* corresponding to *D_total_* = 5 % (Figure 3 D) as threshold for cellular damage. Figure 3 E summaries the definitions for *g_1_*, *g_2_*, *g_3_*, and *g_4_*. It is notable that relations between *g_1_*, *g_2_*, *g_3_*, and *g_4_* may change when altering excitation duration and NP area density.

With the above analysis, the targeted denaturation should meet three criteria simultaneously:

1. A heterogeneous denaturation (*g_1_* < heating power < *g_2_*);
2. Sufficient denaturation at P_1_ (heating power > *g_3_*);
3. Minimal *D_total_* to avoid cellular damage (heating power < *g_4_*).

Bases on these criteria above, we analyzed a large combination of NP arrays and excitations by altering the excitation durations (0.1 ns – 1 ms), NP area densities (0.09 −100 μm^−2^), and heating powers (10^−3^ – 10^9^ μW). First, we generate a 3D map for the boundaries of heterogeneous denaturation. Figure S7 A shows critical surfaces corresponding to *g_1_* and *g_2_*, where the intersection (dotted line) indicates absence of heterogeneous denaturation. Projecting this dotted line in a 2D plot (Figure 4 A) shows the limit for the heterogeneous denaturation window, where targeted denaturation is possible. Next, we investigated boundaries for the second and third criterion located in this window. Figure S7 B&C show critical surfaces of *g_3_* and *g_4_* respectively, where targeted denaturation should between surfaces of *g_3_* and *g_4_* to achieve sufficient denaturation while avoiding cellular damage. Lastly, we combined all the boundaries discussed above and generated a 3D map for targeted protein denaturation (Figure 4 B). A lower limit (maximum of *g_1_* and *g_3_*) and an upper limit (minimum of *g_2_* and *g_4_*) can be set, within which there is targeted denaturation. For example, our results suggest that NCH is established for the case of 10 ns excitation duration at 9 μm^−2^, and target denaturation can be achieved for a heating power within 14 – 81 μW. In contrast, for the case of 1 ms excitation duration at 100 μm^−2^, targeted denaturation does not exist regardless of the heating power since MCH is established. It should be emphasized that these results are based on α-chymotrypsin, while other proteins may have different thresholds. However, we anticipate a similar trend for other proteins when the Arrhenius model is applied. NCH can lead to novel applications in selectively manipulating protein activity by molecular hyperthermia,^[16, 26]^ or membrane permeability,^[27, 28]^ On the other hand, macroscale-collective heating results in uniform heating and widespread protein denaturation for applications such as cancer thermal therapy.^[4, 5]^ Therefore, our analysis provides a framework and guideline to achieve targeted protein denaturation while avoiding widespread damage.

**Figure 4.**
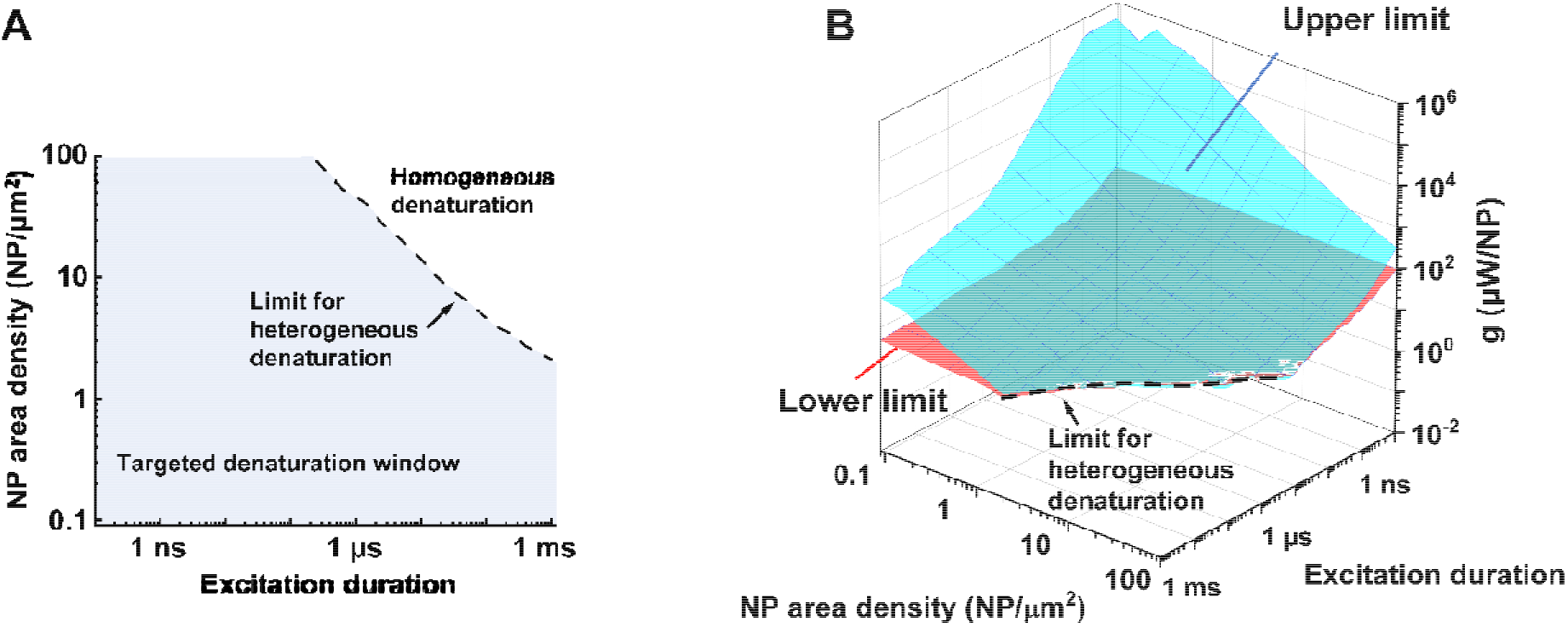
Limits and window for targeted protein denaturation. (A) Projection of the limit for heterogeneous protein denaturation (dotted line), Region below the line (blue area) demonstrated the window for heterogeneous protein denaturation and possible targeted denaturation provided with propriate heating power (shown in B). (B) Lower limit (red surface) and upper limit (blue surface) in terms of excitation duration and NP area density. Region between red surface and blue surface indicates window for targeted protein denaturation.

### 2.3. NCH is insufficient for TRPV1 activation

Lastly, we investigated whether NP array heating could selectively activate TRPV1 channel. TPRV1 channel activation can be treated as a heat-activated chemical reaction described by a two-state model (Equation S6-S9). Previous studies have demonstrated that the TRPV1 channel can be activated within milliseconds at temperatures ranging from 40 – 53 °C (Figure 5 A).^[34, 42]^ Following the framework in Figure 5A, we further investigated TRPV1 activation by NP array heating within 40-53 °C. It is notable that, considering the reversibility for the activation and deactivation of TRPV1 channel,^[34]^ the representative time point for TRPV1 activation analysis is chosen at the end of the excitation duration. Figure 5 B-C illustrates the Δ*T* (TRPV1 normalized activation) profile for case 15-18, with the same NP area density at 9 μm^−2^ and different excitation durations and heating powers (*g*) at 10 ns and 1.69 μW, 10 μs and 0.675 μW, 10 ms and 0.25 μW, 10 s and 0.25 μW respectively. For case 15, NCH is observed due the short excitation duration, but there is no TRPV1 activation. As excitation duration increases, MCH is established (case 16-18), resulting in widespread TRPV1 activation (case 17&18). Here we further quantified the TRPV1 activation by comparing the normalized activation at P_1_ and P_2_ for case 15-18. Figure 5 D shows similar TRPV1 activation at P_1_ and P_2_ for each case, indicating either no TRPV1 activation or widespread TRPV1 activation. We further investigated a large number of conditions by altering excitation duration (1 ms – 1 s), NP area density (0.09 – 100 μm^−2^) and heating power (10^−2^ – 10^2^ μW). Here we set a threshold of on-target TRPV1 activation (at P_1_) at 50 %, and off-target TRPV1 activation (at P_2_) at 10 % to detect any nonspecific excitation.

**Figure 5.**
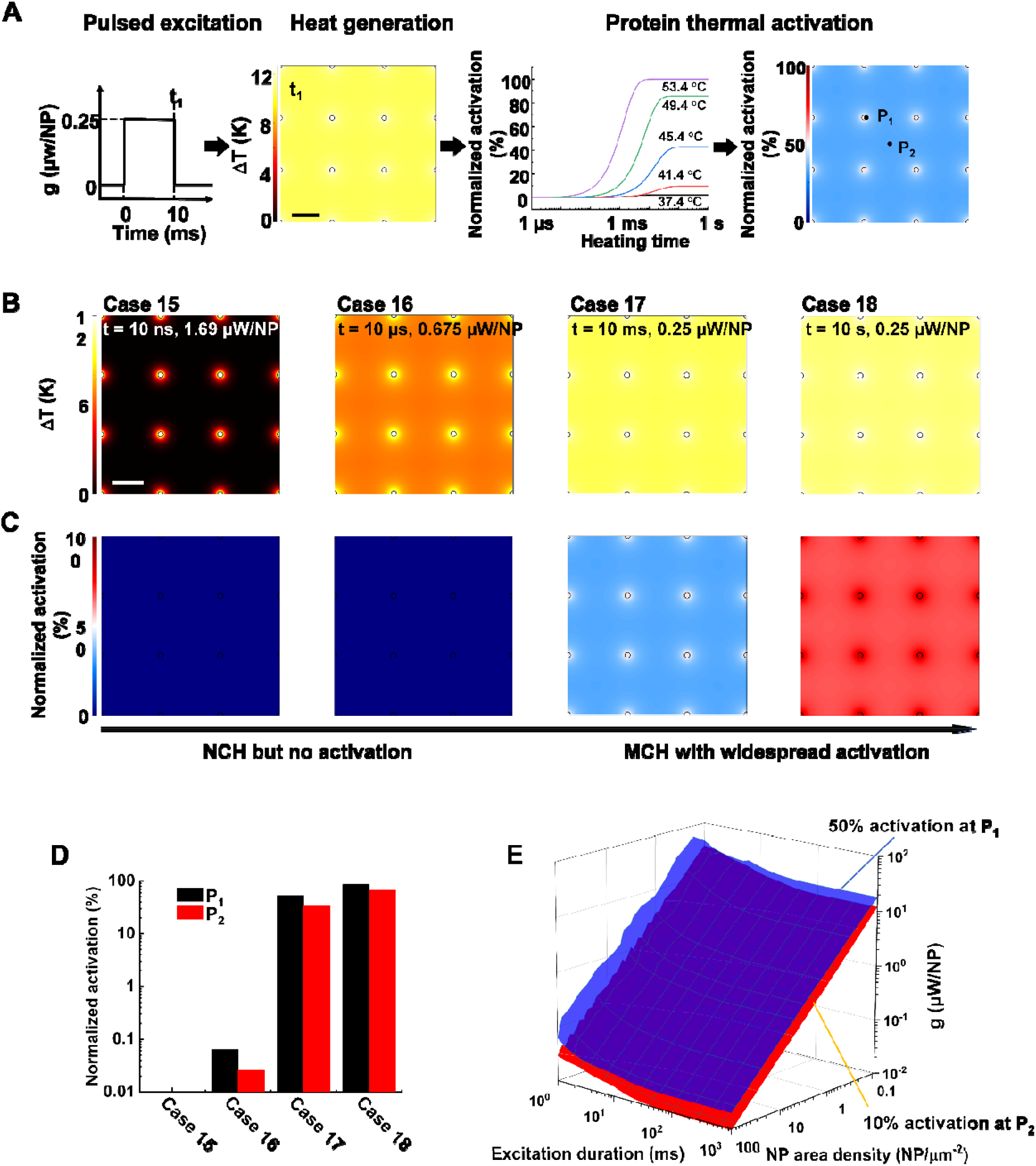
Analysis of TRPV1 channel activation by NP array heating. (A) Schematic of the analysis framework for nanoparticle (NP) array heating and the resulting TRPV1 channel activation. External excitation of the NP array heats up the NP and the surrounding media. The temperature increase drives TRPV1 channel to open, which can be described by two-state model. NP area density: 9 NP/μm^2^, excitation duration: 10 ms, heating power: 0.675 μW/NP. (B) Δ*T* profile and (C) TRPV1 normalized activation profile for cases 15-18. NP area density is 9 NP/μm^2^. Excitation durations and power intensity are shown in the figure. Scalebar represents 200 nm. (D) TRPV1 normalized activation at P_1_ (NP-water interface) and P_2_ (mid-point between particles) for case 15-18. (E) Threshold for on-target TRPV1 activation (50%, at P_1_, blue surface) and off-target TRPV1 activation (10%, at P_2_, red surface).

Figure 5 E shows the threshold for on-target TRPV1 activation (blue surface) is always higher than off-target activation, indicating that on-target TRPV1 activation always comes with off-target activation. Thus, there is always widespread TRPV1 activation by NP array heating because of the mismatch between the short excitation durations (nanosecond timescale) required for NCH and the millisecond timescale needed for channel activation. It should be noted, however, that it is unclear if TRPV1 can be activated at faster rates above 53 °C. According to molecular dynamics simulations of the TRPV1 thermal-gating mechanism, channel opening was not observed within the 200 ns time frame of the simulations at temperatures of 60 and 72 °C, suggesting that TRPV1 activation may be unlikely to occur at nanosecond timescale.^[43]^

## 3. Conclusion

In this study, we investigated how single pulse heating of nanoparticle array affects biological activity, specifically protein thermal denaturation and activation. We found that excitation duration and NP area density are primary factors that determine a window for targeted denaturation. Combined with heating power, we defined quantified boundaries for targeted protein denaturation inside the heterogeneous denaturation window. On the other hand, nanoscale selective activation of the thermally-sensitive ion channel TRPV1 is not feasible based on our current understanding of its millisecond activation kinetics, and nanoparticle array heating leads to widespread TRPV1 activation. This work clearly elucidates physical limits of biological responses driven by NP heating and provides valuable guidance for designing innovative biomedical applications.

## Supporting information

Supplemental information includes Methods, Figures S1 to S7, Tables S1 to S4

## ABBREVIATIONS

NP: nanoparticle;
TRPV1: transient receptor potential cation channel subfamily V member 1;
NCH: nanoscale-selective heating;
MCH: macroscale-collective heating;
SLM: spatial light modulation;
ROS: reactive oxygen species.

## ACKNOWLEDGMENT

The research reported in this work was partially supported by the National Institute of General Medical Sciences (NIGMS) of the National Institutes of Health (award number R35GM133653), the t2019 Collaborative Sciences Award from the American Heart Association (award number 19CSLOI34770004), and the High-Impact/High-Risk Research Award from the Cancer Prevention and Research Institute of Texas (award number RP180846). The content is the sole responsibility of the authors and does not necessarily represent the official views of the funding agencies.

## Notes

### Competing Interest Statement

The authors have declared no competing interest.

