## Supplemental information includes Methods, Figures S1 to S7, Tables S1 to S4 for "Single Pulse Heating of Nanoparticle Array for Biological Applications"

**This PDF file includes:**

Methods

Figures. S1 to S7

Tables S1 to S4

References (1 to 28)

**Methods**

**Analytical solution for single particle heating**

The model for single particle heating consists of a 30 nm spherical nanoparticle surrounded by water, where heat is generated uniformly within the particle and is then diffused into surrounding water. Although some studies suggest that nanoscale thermal diffusion can be different from Fourier’s law,^[1]^ we are mainly concerned with the heating in water with a very small mean free path and similar NP array heating analysis with Fourier’s law has found in well agreement with experimental results.^[2]^ Therefore, we calculated the temperature profile for single NP heating by analytical solution to the heat equation based on Fourier’s law. Here we used 30 nm gold nanoparticles in our models and other NP sizes and materials could be easily modeled under the same framework. The governing equations and the boundary and initial conditions are listed in Table S1. Our results suggest that different NP materials with the same heat generation will have a similar heating profile in water (Figure S2 A), which is the main target for analyzing heating in biological tissues. According to the previous studies, a large temperature increase can occur for NPs that are excited by nanosecond laser pulses and that the surrounding water can superheat well above 100 ºC without phase change.^[3, 4]^ Therefore, we used a heating power (*g*) independent of NP absorption properties to represent the excitation of NPs by different methods, such as laser or magnetic field (Figure S1), and assumed no bubble formation. This heating model was used throughout the remaining of the paper. The analytical solution for this heat transfer model is listed in Table S2^[5]^ and was calculated using Matlab (2019b).

**Multi-particle heating**

For multi-particle heating, we consider the scenario where gold nanoparticles are distributed on the surface of a cell. Because the size of the NP (30 nm) is sufficiently small compared to cells (~10 µm), we neglect the curvature of the cell membrane and simply treat it as a 2D flat plane. The model used to study multiple particle heating is an NP array distributed on a 10 µm × 10 µm plane immersed in water (Figure S2 B). The size of NP arrays was chosen to approximate that of a single cell, and the starting temperature (*T_room_*) was taken as the physiological temperature of 37 °C (310.15 K). For distribution of NPs, we consider three patterns: the square pattern, hexagonal pattern, and the random pattern. The temperature profile was solved by superposition of the analytical solution for single nanoparticle heating:

$$T_{M}\left( \boldsymbol{r},t \right)=\sum_{i=1}^{N} T_{s}^{i}\left( \boldsymbol{r}^{i},t \right)$$

(S1)

where $T_{M}$ is the temperature profile for multi-particle heating and $T_{s}^{i}$ is the analytical solution for *i*^th^ single particle heating (Table S1&2). ***r*** is the position and *t* is time.

We confirmed the accuracy of the superposition method by comparing the temperature profile of the superposition solutions with that of the finite element model (FEM) solutions. First, we developed corresponding FEM models for NP array heating. Figure S2 C illustrates the geometry of the FEM model, where a 5 × 5 NP array (*d_NP_* = 30 nm) is surrounded by water, and the inter-particle gap is 70 nm (NP area density is 100 NP/µm^2^). The water domain is a sphere with a diameter of 1 µm. The boundary of the water domain (sphere surface) is set as a constant temperature (310.15 K) boundary condition. To confirm mesh independence, we compared *∆T_NP_* and *∆T_water_* with different mesh settings (Figure S2 D). Both *∆T_NP_* and *∆T_water_* approach constant values for finer mesh (mesh vertices > 10000), and we adopted a mesh setting with 44068 vertices, which is sufficiently accurate. Next, we confirmed the accuracy of the superposition method by comparing the FEM solution with the superposition solution. Figure S2 E shows the superposition solution and the FEM solution, Point 1 (P_1_) is located at the NP-water interface and Point 2 (P_2_) is located at the mid-point between NPs. The two solutions match well and confirm the accuracy.

We then compared the *∆T_NP_* and *∆T_water_* with different thermal resistance at the NP-water interface. For typical thermal resistance values (10^-7^ – 10^-9^ K∙m^2^∙W^-1^),^[6-14]^ Figure S2 F shows the thermal resistance has a significant effect on *∆T_NP_* whereas it has a negligible effect on *∆T_water_*. Since we are mainly focused on *∆T_water_* profile in this analysis (biological responses in the solution), we ignore the interfacial thermal resistance in our heat transfer model (Table S2). This will give the same temperature for the NP and the water immediately next to the NP surface.

**Random NP array generation and analysis**

For the random pattern, the NP array was generated random coordinate selection on a 10 µm × 10 µm plane. To avoid physical overlapping of NPs, we omitted NPs that overlapped each other and regenerated them to eliminate physical overlap. We generated a large number (10^2^-10^6^) of random NP arrays and calculated the statistical distribution of the inter-particle distances for all arrays (Figure S5 B).

**Protein denaturation**

The protein denaturation was treated as a first-order kinetic model:^[15]^

$$N\underset{\to}{k(T_{m})}D$$

(S2)

where $N$ and $D$ represent the native and denatured states, respectively. The reaction rate was determined by the Arrhenius model:

$$k\left( T_{m} \right)=Ae^{\frac{-E_{a}}{RT_{m}}}$$

(S3)

Normalized denaturation was evaluated by the following equation:

$$D\left( \boldsymbol{r},t \right)=\left( 1-e^{-\int k\left( Tm \right)dt} \right)\times100\%$$

(S4)

We have previously reported the feasibility of α-chymotrypsin photodenaturation by molecular heating.^[16]^ in this work we use α-chymotrypsin as our model protein. The prefactor (*A*) of protein denaturation is 9.75×10^38^ s^-1^ and the activation energy (*E_a_*) is 244.05 kJ∙mol^-1^.^[17]^

Protein denaturation was evaluated by assuming uniform protein distribution on the plane. The total protein denaturation (*D_total_*) was defined as follows:

$$D_{total}=\frac{Denatured protein on the plane}{Total protein on the plane}$$

(S5)

**TRPV1 channel activation**

The TRPV1channel activation and deactivation can be described by the two-state model:^[18, 19]^

|  | $k_{on}$  $C⇆O$  $k_{off}$ | (S6) |
| --- | --- | --- |

*C* represents the closed state of the channel, and *O* represents the open state of the channel. The steady-state transmembrane current is dependent on temperature, which can be described by Boltzmann equation (Table S4):

$$I_{ss}=I_{l}e^{\frac{-{\Delta H}_{l}}{RT}}+\frac{I_{max}e^{\frac{-\Delta H}{RT}}}{1+e^{\frac{-\Delta H-T\Delta S}{RT}}}$$

(S7)

The transient transmembrane current can be calculated by the following equation (Table S4):

|  | $I=I_{ss}\left( 1-e^{-\int kdt} \right)$ | (S8) |
| --- | --- | --- |
| where |  |  |
|  | $k=Ae^{\frac{-\Delta H-T\Delta S}{RT}}$ | (S9) |

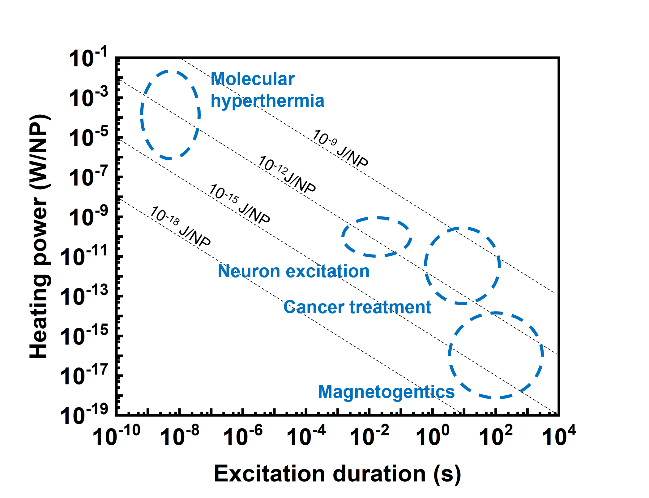

**Figure S1. Power-duration map.** Dashed circles indicate approximate locations of molecular hyperthermia,^[20,21]^ neuron excitation,^[22,23]^ cancer treatment,^[24,25]^ and magnetogentics.^[26,27]^ Dotted black lines indicate equal amounts of input energy per nanoparticle.

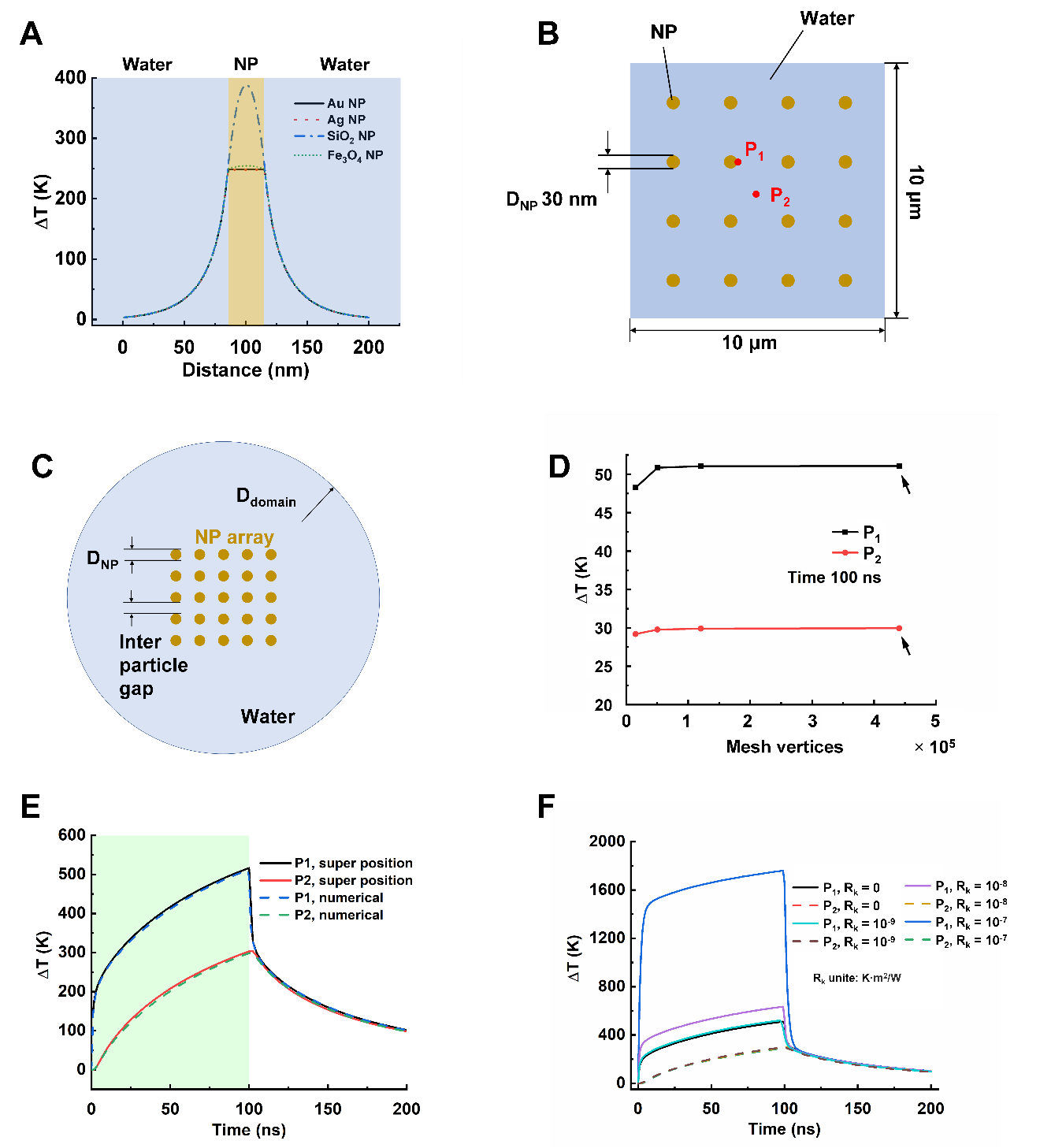

**Figure S2. Model set up.** (A) *∆T* profile for different NP materials. D_NP_ = 30 nm, heating power per NP (*g*) is 35.6 μW, excitation duration 10 ns, time at 10 ns. *∆T* profile in NP may change due to different thermal properties, whereas the *∆T* profile are almost identical in water. (B) Schematic for the superposition model. The blue square represents the domain of the model, gold dotes represents NP. The size of domain is 10 μm ×10 μm and the size of NP is 30 nm. The 2D NP array is immersed in water. P_1_ is in NP and P_2_ is at the mid-point between NPs. (C) Schematic for the finite element model (FEM), *d_domain_* = 1 µm, *d_NP_* = 30 nm, and inter-particle gap = 70 nm, NP area density = 100 µm^-2^. (D) *∆T* with different mesh settings (time at 100 ns). The arrows indicate setting used in further analysis. (E) Validation of the superposition method by comparing the superposition solution and the FEM solution. (F) *∆T*(*t*) profile with various NP-water interfacial thermal resistance. While the interfacial thermal resistance changes the NP temperature, but it has negligible effect on *∆T* profile in water.

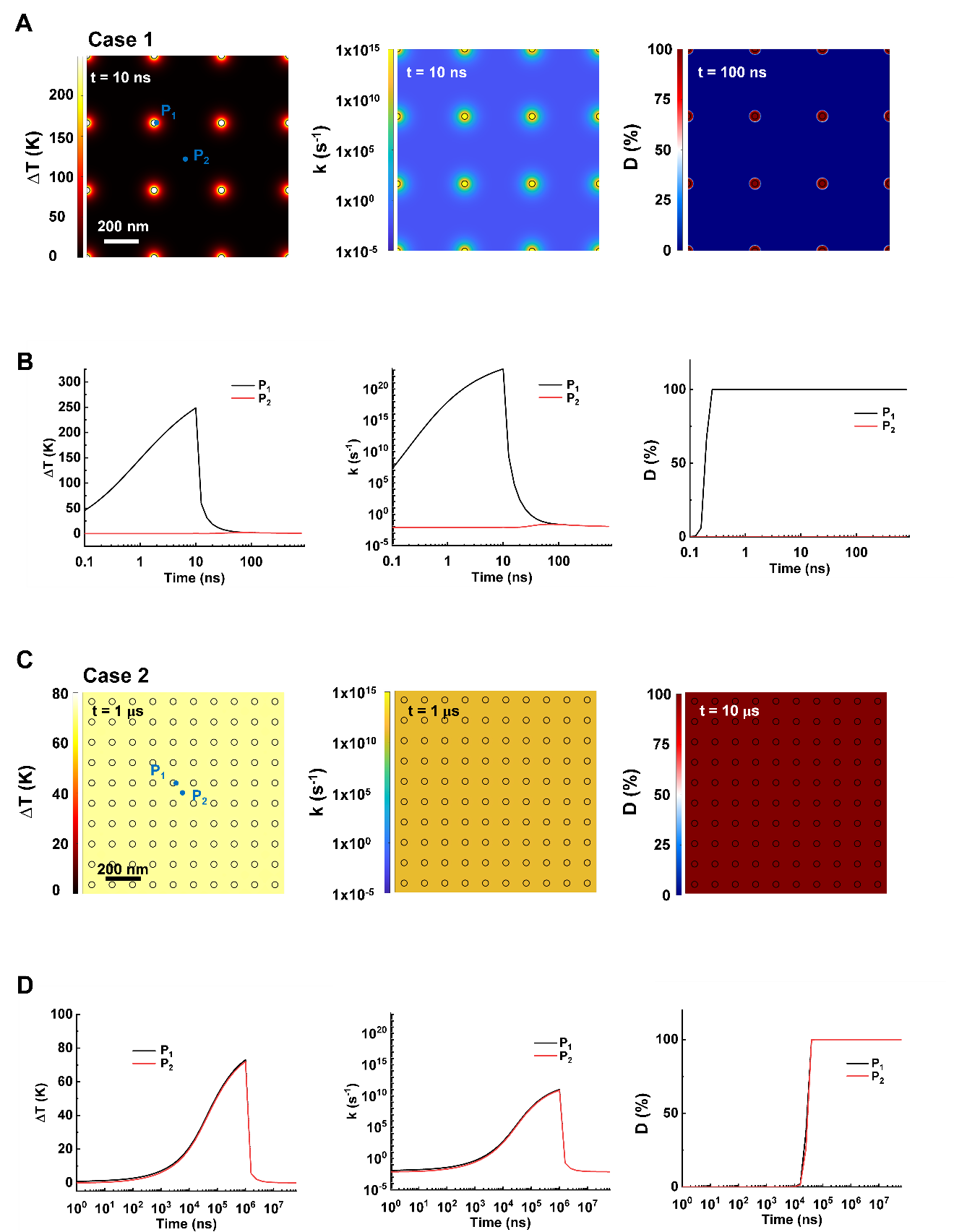

**Figure S3. *∆T*, reaction rate and protein denaturation profile (*D*) for case 1 (A&B) and case 2 (C&D).** (A&B) Nanoscale confined heating (NCH), model size 10 μm ×10 μm, *d_NP_* =30 nm, *g* = 35.6 μW, NP area density 9 µm^-2^, excitation duration 10 ns. (C&D) macroscale collective heating (MCH), model size 10 μm ×10 μm, *d_NP_* = 30 nm, *g* = 18 nW, NP area density:100 µm^-2^, excitation duration: 1 µs.

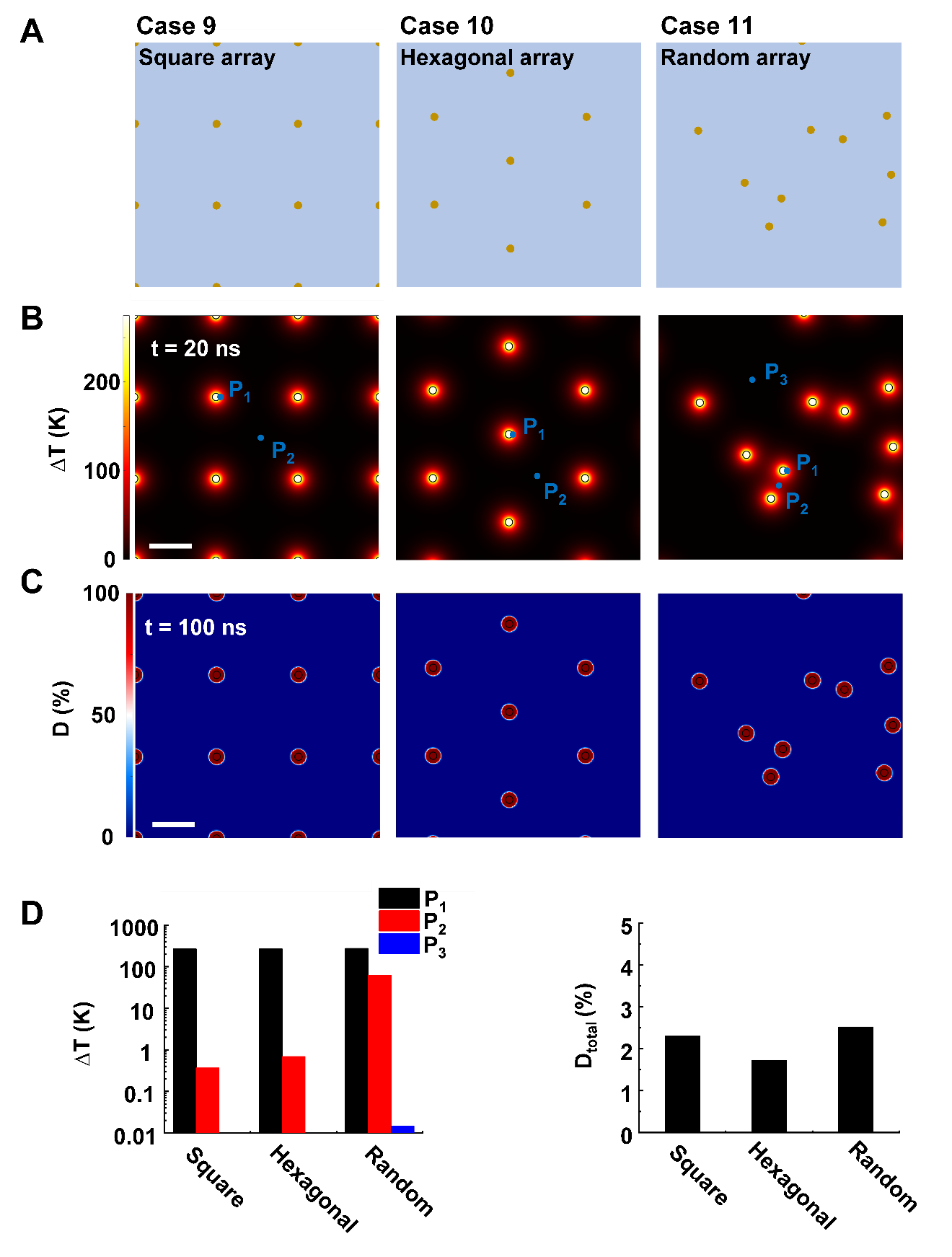

**Figure S4. Effect of NP distribution on heating and protein denaturation.** (A) Schematic of NP distribution for case 9-11, square (case 9) hexagonal (case 10) random (case 11). (B) *∆T* profile and (C) *D* profile for cases 9-11. (D) *∆T* at representative locations (P_1_: NP-water interface, P_2_: mid-point between NPs, for random distribution, P_2_ is inside a NP cluster, P_3_ is outside the NP clusters) and *D_total_* for case 9-11. For all cases, NP area density is 9 µm^-2^, *g =* 35.6 µW. Scalebar represents 200 nm.

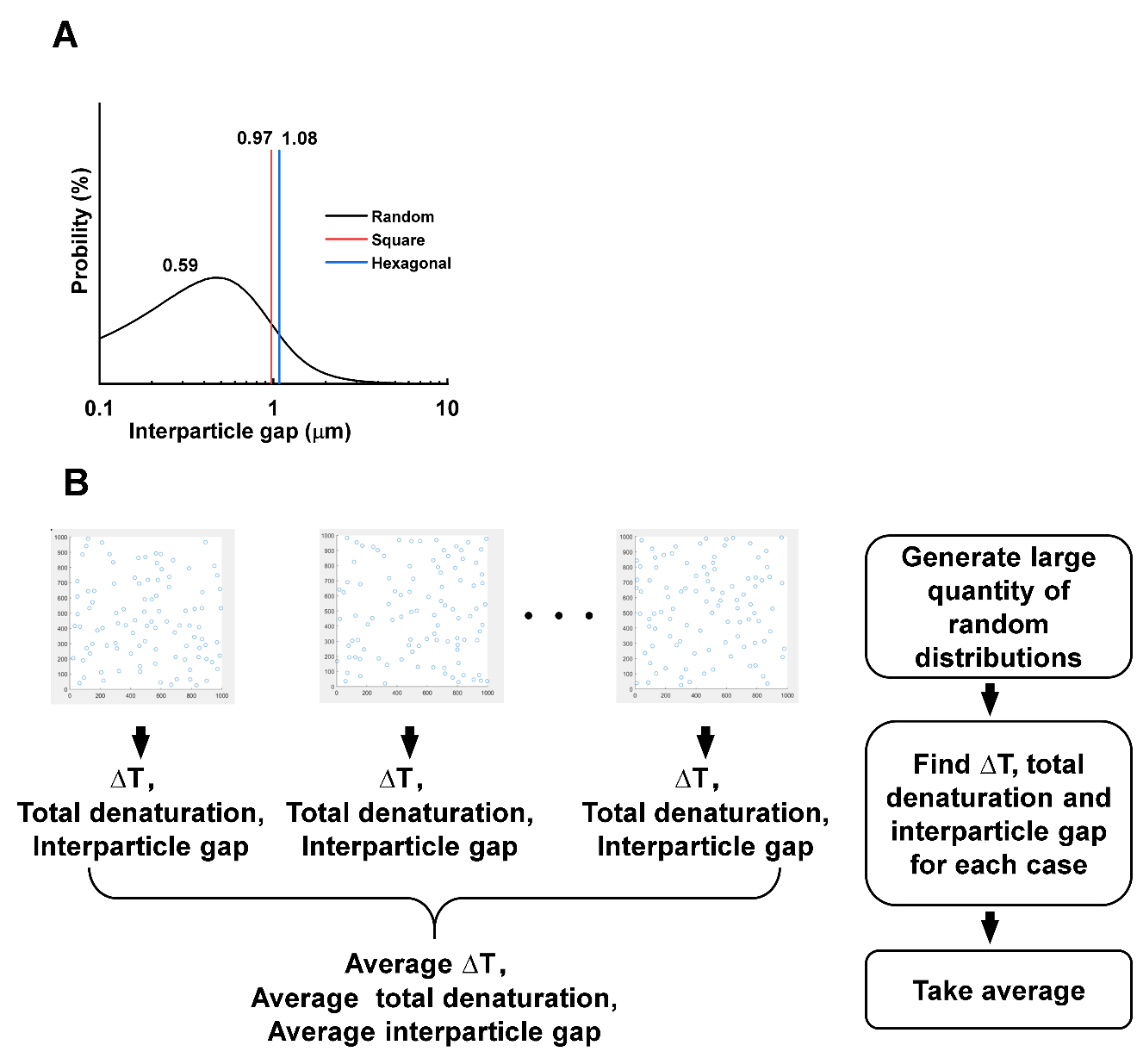

**Figure S5. Analysis of the random NP distribution and inter-particle distance**. (A) Inter-particle gap distributions for random, square, and hexagonal distributions The NP area density is 1 µm^-2^. (B) Schematic illustration for the random pattern distribution analysis.

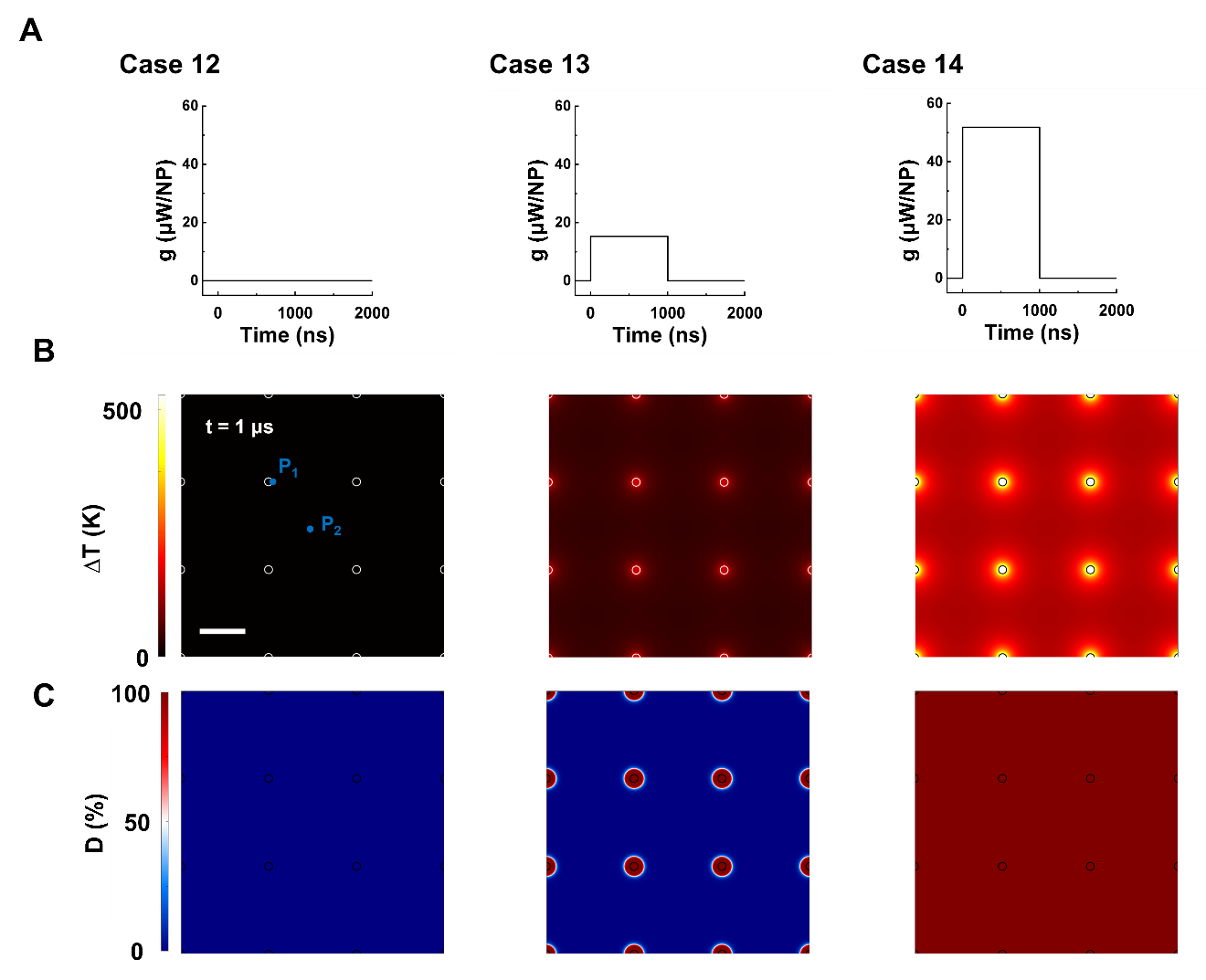

**Figure S6. *∆T* and *D* profile for case 12-14.** Simulation condition: Excitation duration 1 μs, NP area density 9 µm^-2^. (A)Schematic of *g* for case 12-14, 0.002 μW (case 12), 15.3 μW (case 13), and 51.8 μW (case 14). (B) *∆T* profile and (C) *D* profile for cases 12-14, scale bar represents 200 nm.

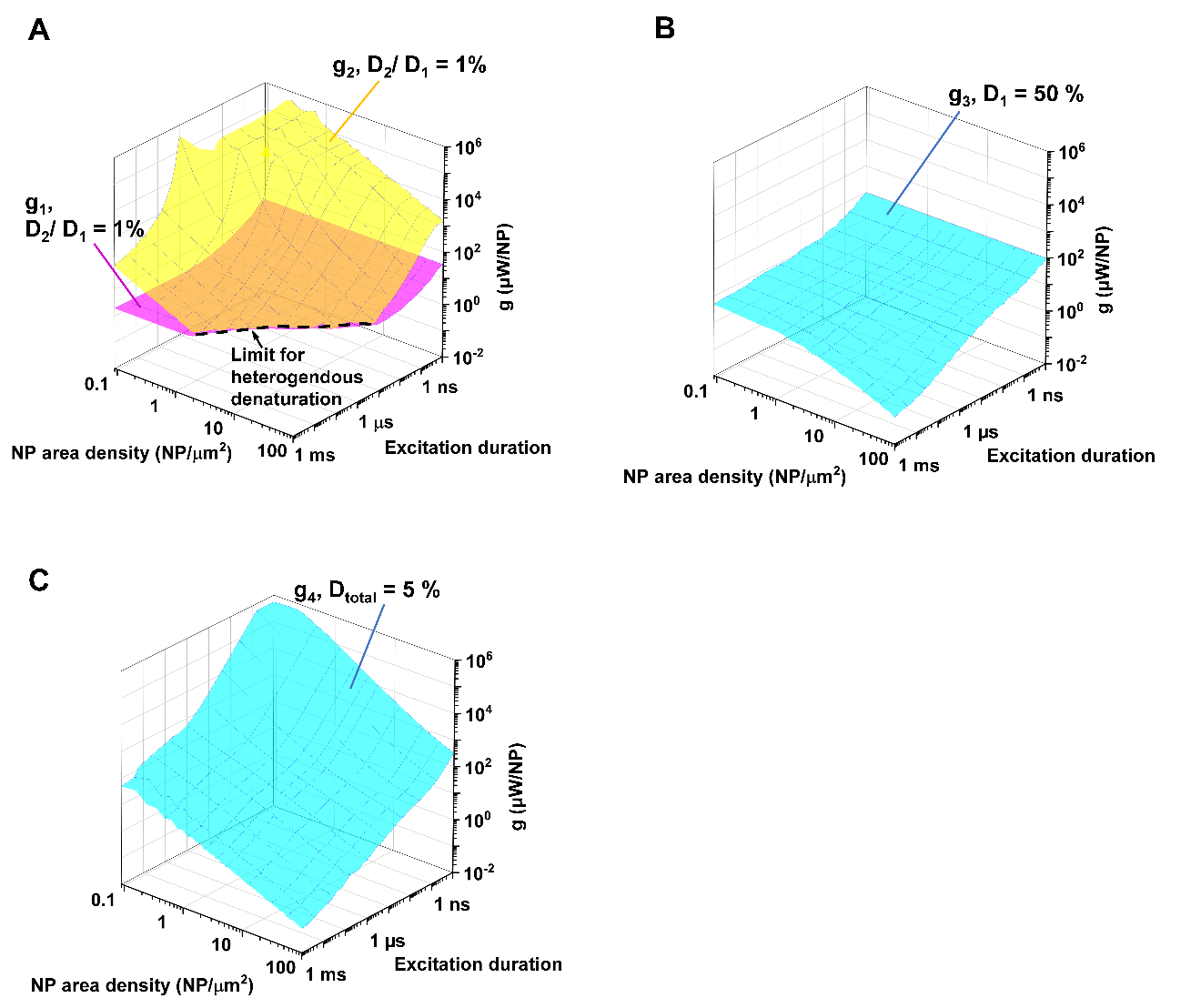

**Figure S7. Limits and window for targeted denaturation sand targeted denaturation.** (A) *g_1_* (purple surface) and *g_2_* (yellow surface) in terms of excitation duration and NP area density. Region between purple surface and yellow surface indicates window for heterogeneous protein denaturation, with no denaturation below *g_1_* and widespread denaturation above *g_2_*. The position where the two surfaces intersect suggests diminishing window for heterogeneous protein denaturation (dotted line). (B) *g_3_* (*D_1_* = 50%) in terms of the excitation duration and NP area density. Region under this surface indicates insufficient denaturation (*D_1_* < 50%). (C) *g_4_* (*D_total_* = 5 %) in terms of the excitation duration and NP area density. Region above this surface indicates cellular damage.

**Table S1. Governing equations and boundary and initial conditions for single nanoparticle heating.**

| Discription | Equations | Equation number |
| --- | --- | --- |
| Governing  equation | $\frac{1}{\alpha_{Au}}\frac{\partial T_{Au}}{\partial t}=\frac{1}{r^{2}}\frac{\partial}{\partial r}\left( r^{2}\frac{\partial T_{Au}}{\partial r} \right)+\frac{G_{v}}{k_{Au}}g=G_{v}\cdot V_{NP}$ | (11) |
|  | $\frac{1}{\alpha_{water}}\frac{\partial T_{water}}{\partial t}=\frac{1}{r^{2}}\frac{\partial}{\partial r}\left( r^{2}\frac{\partial T_{water}}{\partial r} \right)$ | (12) |
| Boundary conditions | $-k_{Au}\left. \frac{\partial T_{Au}}{\partial r} \right\vert_{r=R_{NP}}=-k_{water}\left. \frac{\partial T_{water}}{\partial r} \right\vert_{r=R_{NP}}$ | (13) |
|  | $\left. T_{Au} \right\vert_{r=R_{NP}}=\left. T_{water} \right\vert_{r=R_{NP}}$ | (14) |
|  | $\left. \frac{\partial T}{\partial r} \right\vert_{r=0}=0$ | (15) |
|  | $\left. T \right\vert_{r=\infty}=T_{ref}=310.15 K$ | (16) |
| Initial condition | $\left. T \right\vert_{t=0}=T_{ref}=310.15 K$ | (17) |
| Volumatic heat source | $G_{v}=\frac{g}{V_{NP}}$ | (18) |

**Table S2. Analytical solution for single nanoparticle heated by a rectangular pulse.^[28]^**

| Heating process | $\left. T_{Au}(r,t) \right\vert_{t<2\mu}= T_{ref}+\frac{{R_{NP}}^{2}}{k_{Au}}\frac{g}{V_{NP}2\mu}\left\{ \frac{1}{3}\frac{k_{Au}}{k_{water}}+\frac{1}{6}\left( 1-\frac{r^{2}}{{R_{NP}}^{2}} \right)-\frac{2R_{NP}b}{r\pi}\int_{0}^{\infty} \frac{exp(-\frac{y^{2}t}{\gamma_{1}})}{y^{2}}\frac{\left( \sin y-y\cos y \right)sin( \frac{ry}{R_{NP}})}{[\left( c\sin y-y\cos y \right)^{2}+b^{2}y^{2}{sin}^{2}y]}dy \right\}$ | (19) |
| --- | --- | --- |
|  | $\left. T_{Au}(r=0,t) \right\vert_{t<2\mu}= T_{ref}+\frac{{R_{NP}}^{2}}{k_{Au}}\frac{g}{V_{NP}2\mu}\left\{ \frac{1}{3}\frac{k_{Au}}{k_{water}}+\frac{1}{6}-\frac{2b}{\pi}\int_{0}^{\infty} \frac{exp(-\frac{y^{2}t}{\gamma_{1}})}{y^{2}}\frac{\left( \sin y-y\cos y \right)}{[\left( c\sin y-y\cos y \right)^{2}+b^{2}y^{2}{sin}^{2}y]}dy \right\}$ | (20) |
|  | $\left. T_{water}(r,t) \right\vert_{t<2\mu}= T_{ref}+\frac{{R_{NP}}^{3}}{rk_{Au}}\frac{g}{V_{NP}2\mu}\left\{ \frac{1}{3}\frac{k_{Au}}{k_{water}}-\frac{2}{\pi}\int_{0}^{\infty} \frac{exp(-\frac{y^{2}t}{\gamma_{1}})}{y^{3}}\frac{\left( \sin y-y\cos y \right)\left[ by\sin y\cos\sigma y-\left( c\sin y-y\cos y \right)\sin\sigma y \right]}{[\left( c\sin y-y\cos y \right)^{2}+b^{2}y^{2}{sin}^{2}y]}dy \right\}$ | (21) |
| Cooling process | $\left. T_{Au}(r,t) \right\vert_{t\geq2\mu}=T_{ref}+\frac{2{R_{NP}}^{3}b}{r\pi k_{Au}}\frac{g}{V_{NP}2\mu}\int_{0}^{\infty} \frac{\exp\left[ -\frac{y^{2}\left( t-2\mu\right)}{\gamma_{1}} \right]-\exp(\frac{-y^{2}t}{\gamma_{1}})}{y^{2}}\frac{\left( \sin y-y\cos y \right)sin( \frac{ry}{R_{NP}})}{[\left( c\sin y-y\cos y \right)^{2}+b^{2}y^{2}{sin}^{2}y]}dy$ | (22) |
|  | $\left. T_{Au}(r=0,t) \right\vert_{t\geq2\mu}=T_{ref}+\frac{2{R_{NP}}^{2}b}{\pi k_{Au}}\frac{g}{V_{NP}2\mu}\int_{0}^{\infty} \frac{\exp\left[ -\frac{y^{2}\left( t-2\mu\right)}{\gamma_{1}} \right]-\exp(\frac{-y^{2}t}{\gamma_{1}})}{y^{2}}\frac{\left( \sin y-y\cos y \right)}{[\left( c\sin y-y\cos y \right)^{2}+b^{2}y^{2}{sin}^{2}y]}dy$ | (23) |
|  | $\left. T_{water}(r,t) \right\vert_{t\geq2\mu}= T_{ref}+\frac{{R_{NP}}^{3}}{rk_{Au}}\frac{g}{V_{NP}2\mu}\left\{ -\frac{2}{\pi}\int_{0}^{\infty} \frac{\exp\left[ -\frac{y^{2}\left( t-2\mu\right)}{\gamma_{1}} \right]-\exp(\frac{{-y}^{2}t}{\gamma_{1}})}{y^{3}}\frac{\left( \sin y-y\cos y \right)\left[ by\sin y\cos\sigma y-\left( c\sin y-y\cos y \right)\sin\sigma y \right]}{[\left( c\sin y-y\cos y \right)^{2}+b^{2}y^{2}{sin}^{2}y]}dy \right\}$ | (24) |
|  | $b= \frac{k_{water}}{k_{gold}}\sqrt{\frac{\alpha_{Au}}{\alpha_{water}}}, c=1-\frac{k_{water}}{k_{gold}}, \gamma_{1}=\frac{{R_{NP}}^{2}}{\alpha_{gold}}, \sigma= \left( \frac{r}{R_{NP}}-1 \right)\sqrt{\frac{\alpha_{gold}}{\alpha_{water}}}$ | (25) |

**Table S3. Parameters for the nanoparticle array heating cases.**

| Case | Excitation duration [ns] | NP area density [µm^-2^] | NP distribution | Heating power per NP (*g*) [µW] | Temperature raise | | | Protein denaturation | |
| --- | --- | --- | --- | --- | --- | --- | --- | --- | --- |
|  |  |  |  |  | *∆T_1_* [K] | *∆T_2_* [K] | *∆T_2_/∆T_1_* [%] | *D_total_* [%] | *D_2_/ D_1_* [%] |
| 3 | 10 | 9 | Square | 35.6 | 248.7 | 0.2 | 0.0 | 2.2 | 0.0 |
| 4 | 100 |  |  |  | 296.8 | 14.0 | 4.7 | 5.4 | 0.0 |
| 5 | 1,000 |  |  |  | 375.8 | 92.9 | 33.7 | 100.0 | 100.0 |
| 6 | 20 | 2.6 |  |  | 267.2 | 0.05 | 0.0 | 0.8 | 0.0 |
| 7 |  | 25 |  |  | 268.5 | 9.6 | 3.6 | 8.3 | 0.0 |
| 8 |  | 100 |  |  | 318.8 | 109.5 | 34.2 | 99.9 | 100.0 |
| 9 |  | 9 | Square |  | 267.3 | 0.4 | 0.0 | 2.3 | 0.0 |
| 10 |  |  | Hexagon |  | 267.8 | 0.7 | 0.0 | 1.7 | 0.0 |
| 11 |  |  | Random |  | 274.6 | 61.8 |  | 2.5 |  |
| 12 |  |  | Square | 0.002 | 2.1 | 0.5 | 24.74 | 0.05 | 71.2 |
| 13 |  |  |  | 15.3 | 162.0 | 40.1 |  | 4.3 | 0.0 |
| 14 |  |  |  | 51.8 | 546.6 | 135.2 |  | 8.1 | 100.0 |

**Table S4. Parameters for the TRPV1 channel activation model.^[18]^**

| Leakage current | $I_{l}$ | 36.2 [nA] |
| --- | --- | --- |
| Enthalpy for leakage current | $\Delta H_{l}$ | 17.0 [kJ∙mol^-1^] |
| Maximum current | $I_{max}$ | 6.8×10^69^ [nA] |
| Activation enthalpy | $\Delta H$ | 422.8 [kJ∙mol^-1^] |
| Transition entropy | $\Delta S$ | 1.3 [kJ∙mol^-1^∙K^-1^] |
